# Human Cytomegalovirus Drives WDR5 to the Virion Assembly Compartment to Facilitate Virion Assembly

**DOI:** 10.1101/2020.08.03.235630

**Authors:** Bo Yang, YongXuan Yao, Hui Wu, Hong Yang, Xue-Hui Ma, Dong Li, Xian-Zhang Wang, Sheng-Nan Huang, Xuan Jiang, Shuang Cheng, Jin-Yan Sun, Zhen-Li Huang, CongJian Zhao, Michael A. McVoy, Jin-Hyun Ahn, Wen-Bo Zeng, Sitang Gong, William J. Britt, Min-Hua Luo

## Abstract

We previously reported that human cytomegalovirus (HCMV) utilizes the cellular protein WDR5 to facilitate capsid nuclear egress. Here, we further show that HCMV infection drives WDR5 to the perinuclear region by a mechanism that requires viral replication and intact microtubules. WDR5 accumulated in the virion assembly compartment (vAC) and co-localized with vAC markers of gamma-tubulin (γ-tubulin), early endosomes, and viral vAC marker proteins pp65, pp28, and glycoprotein B (gB). WDR5 interacted with multiple virion proteins, including MCP, pp150, pp65, pIRS1, and pTRS1, which may explain the increasing WDR5 accumulation in the vAC during infection. WDR5 was then incorporated into HCMV virions and localized to the tegument layer, as demonstrated by fractionation and immune-gold electron microscopy. Thus, WDR5 is driven to the vAC and incorporated into virions, suggesting that WDR5 facilitates HCMV replication at later stage of virion assembly besides the capsid nuclear egress stage. These data highlight that WDR5 is a potential target for antiviral therapy.

**Importance:** Human cytomegalovirus (HCMV) has a large (~235-kb) genome that contains over 170 ORFs and exploits numerous cellular factors to facilitate its replication. In the late phase of HCMV infection cytoplasmic membranes are profoundly reconfigured to establish the virion assembly compartment (vAC), which is important for efficient assembly of progeny virions. We previously reported that WDR5 promotes HCMV nuclear egress. Here, we show that WDR5 is further driven to the vAC and incorporated into virions, perhaps to facilitate efficient virion maturation. This work identified potential roles for WDR5 in HCMV replication in the cytoplasmic stages of virion assembly. Taken together, WDR5 plays a critical role in HCMV capsid nuclear egress and is important for virion assembly, and thus is a potential target for antiviral treatment of HCMV-associated diseases.

## Introduction

Human cytomegalovirus (HCMV) is a ubiquitous pathogen that is highly adapted to its human host. Approximately 50 to 90% of adults are infected globally (1) and in China seroprevalence is as high as 93.7% (2). About 90% of primary HCMV infections in immunocompetent individuals are either asymptomatic or mildly symptomatic. In addition, HCMV is a leading cause of congenital infections worldwide. Following primary infection, the virus remains latent and establishes life-long persistence in its host. However, HCMV can be reactivated and cause severe disease in immunosuppressed or immunodeficient individuals, such as recipients of solid organ or bone marrow transplants and AIDS patients (3–7). In these patients, HCMV infection contributes to multiple end-organ diseases, including pneumonia, gastrointestinal diseases, retinitis, central nervous system diseases, poor engraftment, severe graft-versus-host reaction, and can result in high mortality rates (8–11). In patients receiving hematopoietic stem cell transplantation, the incidence of HCMV disease is as high as 30-70% (12). In addition, congenital HCMV infection is a leading cause of birth defects and is associated with neonatal mortality and permanent developmental sequelae (13–16).

A characteristic feature of HCMV infection is profound remodeling of the secretory and endocytic system (Golgi, trans-Golgi network, and early endosomes) to form nested, cylindrical layers in a structure termed the virion assembly compartment (vAC) (4, 17–22), which is unique in HCMV infection when compared to other herpesviruses (23). Although not essential for virion production, vAC formation has been shown to be necessary for efficient HCMV virion assembly and maturation. Multiple viral and cellular proteins are reported to participate in vAC formation. Viral protein pUL136 is important for vAC formation and efficient virion assembly. Virus mutants lacking UL136 produce less properly enveloped virions along with a larger number of dense bodies compared to wildtype virus (24). Other viral proteins pUL48, pUL94, and pUL103 are also important for vAC development (21). Interestingly, HCMV-encoded miRNAs UL112-1, US5-1, and US5-2 target host secretory pathway factors to facilitate vAC formation (25). Cellular proteins, including RhoB (22), IFITMs (23), STX5 (26), dynein (27), BiP (27, 28), Bicaudal D1 (29), Rab11 and FIP4 (30), have also been shown to localize to vAC.

A number of cellular proteins are relocated to the vAC and subsequently incorporated into virions. Proteomic analyses have identified over 70 viral and cellular proteins in HCMV virions. Virion-associated cellular proteins are involved in a wide range of processes and functions, including ATP binding, Ca^2+^ signaling, signal transduction, transcription, and translation (31). WD repeat-containing protein 5 (WDR5) is a highly conserved WD-40 repeat protein that is critical for regulating multiple cellular processes. The most common function of WD-repeat proteins is to serve as a rigid scaffold for protein-protein interactions, which facilitates formation and/or stabilization of multi-protein complexes (32), such as MOF (males absent on the first)-containing NSL (33), KANSL1 and KANSL2 (33), and histone H3 at lysine 4 (H3K4) methyltransferases of the SET1-family (Set1A, Set1B, MLL1, MLL2, MLL3, and MLL4) (34–37). Thus, previous studies of WDR5 have focused on epigenetics (38–41); however, WDR5 can also stabilize actin architecture to promote multi-ciliated cell formation (42) and is important for embryonic stem cell reprogramming, self-renewal, and maintenance of pluripotency (43, 44).

The role of WDR5 in RNA virus infections has also been investigated. Upon Sendai virus infection, WDR5 is translocated from the nucleus to the mitochondria and is essential for formation of the VISA-associated complex (45). Measles virus recruits WDR5 into viral inclusion bodies in the cytoplasm to facilitate virus replication (46). Our previous work showed that WDR5 promotes HCMV replication through facilitating capsid nuclear egress (47). In the present study, we further demonstrate that in HCMV-infected cells WDR5 protein levels increase in the cytoplasm and that WDR5 is translocated to and accumulates in the vAC. The accumulated WDR5 is incorporated into and localizes within the tegument layer of virions which facilitates virion morphogenesis.

## Results

### Perinuclear accumulation of WDR5 is induced by HCMV infection

It has been reported that cellular protein WDR5 is translocated from the nucleus to mitochondria during Sendai virus infection and to viral replication centers during measles virus infection. As both are RNA viruses (45, 46), we sought to investigate whether a corresponding WDR5 relocation occurs during HCMV infection. We initially determined if WDR5 is redistributed during HCMV infection of immortalized human embryonic lung fibroblasts (HELFs). In mock-infected HELFs, WDR5 had diffuse nuclear localization with a single small perinuclear punctum which coincided with MTOC, whereas in HCMV-infected cells WDR5 was clearly localized in a perinuclear region (Fig 1A). To exclude the possibility that WDR5 perinuclear accumulation is limited to HELFs or infection by HCMV strain Towne, similar experiments were performed with primary cell HELs and U251 glioma cells infected with HCMV strains Towne or AD169, and similar perinuclear WDR5 localization was observed (data not shown). In addition, similar perinuclear localization was also observed during murine cytomegalovirus (MCMV) infection of mouse NIH3T3 fibroblasts (Fig 1B). Interestingly, this effect was not observed during infection of HELFs by other DNA viruses that have not been reported to form vAC-like structures during infection, including herpes simplex virus type 1 (HSV-1), varicella-zoster virus (VZV), adenovirus (AdV), or autographa californica multiple nuclear polyhedrosis virus (AcMNPV) (Fig 1C). These data suggest that the perinuclear accumulation of WDR5 is specifically induced by HCMV or MCMV infection.

**Fig 1.**
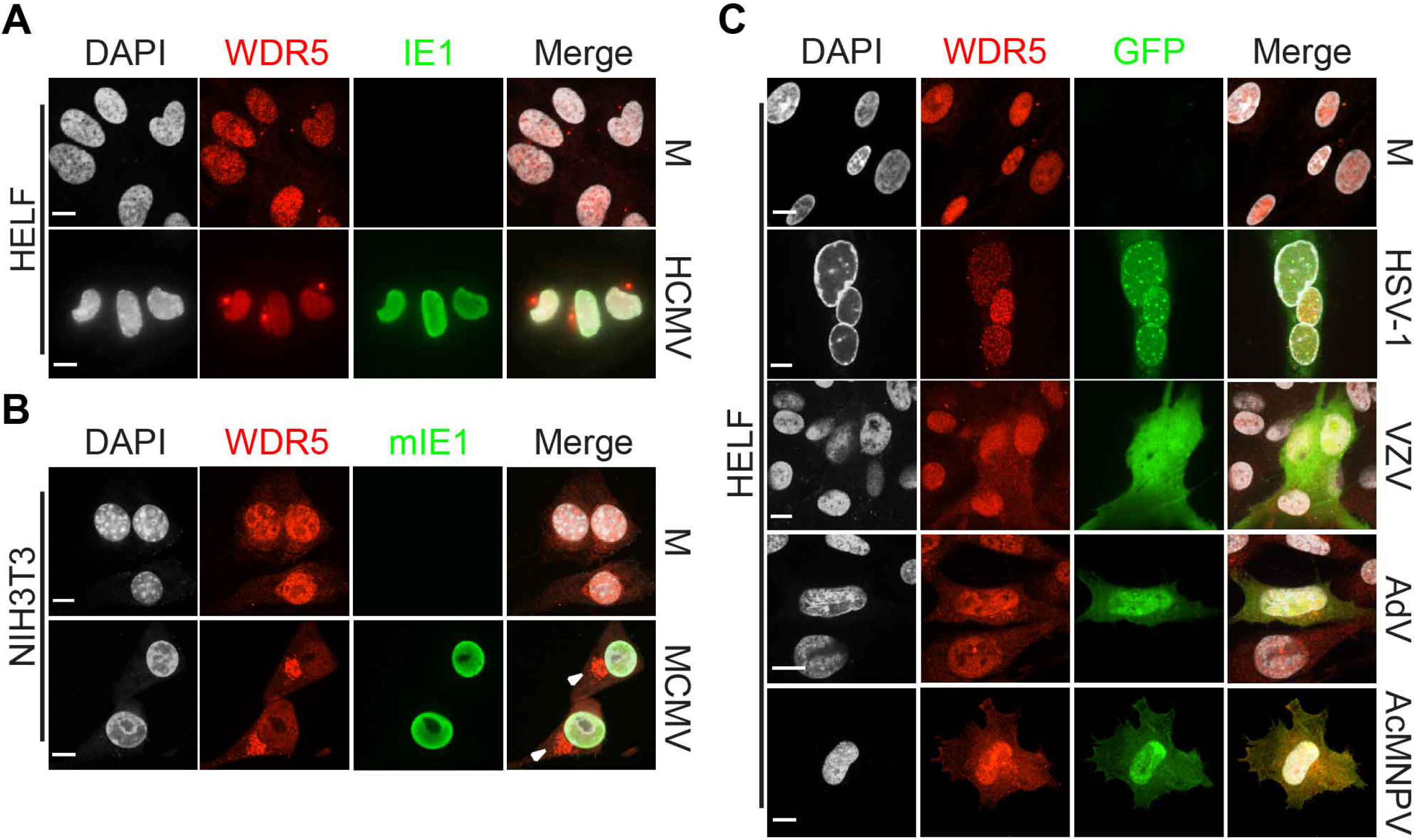
Perinuclear relocalization of WDR5 is specifically induced by HCMV and MCMV infection. HELFs (A and C) or NIH3T3 (B) were infected with the indicated viruses at an MOI of 1. At 48 hpi, cells were fixed and stained with antibodies against WDR5(red) and with either HCMV IE1 protein (A) (green) or MCMV mIE1 (green) (B). Infected cells (C) were detected using corresponding GFP-expressing recombinant viruses. All antibodies used in IFA were mouse monoclonal antibodies. Nuclei were counterstained with DAPI (white). M, mock-infected cells; scale bar = 10 μm.

### WDR5 perinuclear accumulation progresses with HCMV infection

To gain further insights into changes in subcellular localization of WDR5 during HCMV infection, a time course confocal microscopy study was performed from 12 to 96 hpi. Focal perinuclear accumulation of WDR5 was first observed at 24 hpi and increased as the infection progressed (Fig 2A and B).

**Fig 2.**
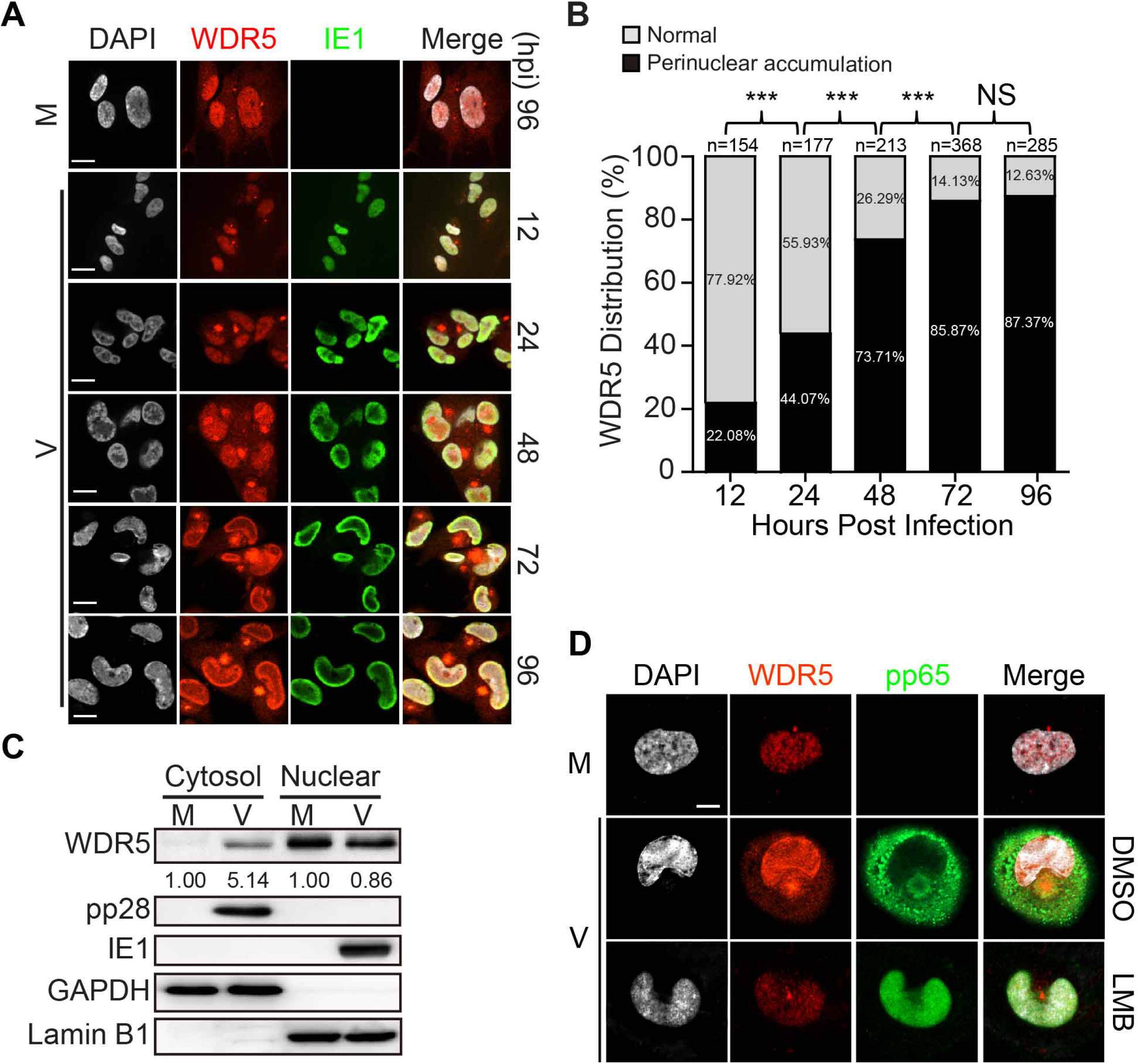
HCMV infection induces re-localization and oligomerization of WDR5. (A) HELFs were mock- (M) or HCMV-infected (V) at an MOI of 3 then processed for immunofluorescence with antibodies against IE1 (green), WDR5 (red), or nuclei (DAPI, white) at the indicated times post infection. Images shown were representative of three independent experiments. All antibodies used in IFA were mouse monoclonal antibodies. Scale bar = 10 μm. (B) Over 100 cells in five representative fields from each independent experiment described in (A) were counted to determine the percentages of infected cells exhibiting diffuse nuclear localization of WDR5 (normal) versus accumulation of WDR5 in perinuclear foci. Data were analyzed using the chi-squared test. N, number counted cells; NS, not significant; ***, P < 0.001. (C) HELFs were mock- (M) or HCMV-infected (V) at an MOI of 3. At 96 hpi cytosolic and nuclear fractions were prepared and analyzed by IB for WDR5, pp28, IE1, GAPDH, or Lamin B1. (D) HELFs were mock- or HCMV-infected at an MOI = 1. At 48 hpi, infected cells were treated with DMSO or nuclear export inhibitor Leptomycin B (LMB, 5μM) for 12 h, and were co-stained for WDR5 (red), viral protein pp65 (green) and nuclei were counterstained with DAPI (white). All antibodies used in IFA were mouse monoclonal antibodies. Scale bar = 10 μm.

To further investigate the impact of HCMV infection on WDR5 and its cytosolic/nuclear distribution, HCMV- and mock-infected cells were harvested at 96 hpi and cytosolic and nuclear fractions were analyzed by immunoblot (IB). The efficiency of nuclear and cytosolic fractionation was confirmed by probing for cellular markers GAPDH and Lamin B1, which localize to cytoplasm and nuclei, respectively, as well as for viral proteins pp28 (pUL99) and IE1, which also localize to the cytosol and nucleus, respectively (Fig 2C). In mock-infected cells, WDR5 was located in the nuclear fraction and at a much lower level in the cytosol. In contrast, HCMV-infected cells contained less WDR5 protein in the nuclear fraction, while with a significantly higher level of WDR5 protein in the cytosolic fraction (Fig 2C). Previous studies have shown that nuclear membranes are significantly remodeled during HCMV infection (27). To exclude a possibility that higher cytosolic protein level of WDR5 was a result from simple diffusion, leptomycin B (LMB) was used to inhibit nuclear export. Nuclear retention of WDR5 was clearly observed and perinuclear accumulation of WDR5 was prevented by LMB treatment (Fig 2D). These data suggested that nuclear WDR5 relocated to the cytoplasm during HCMV infection is not due to diffusion.

### WDR5 perinuclear accumulation requires viral replication

To determine the kinetic class of viral gene expression required to alter WDR5 localization, HELFs were incubated with HCMV virions that were UV-irradiated to block all *de novo* viral gene expression or with infectious virions in the presence of the viral DNA polymerase inhibitors ganciclovir (GCV) or phosphonoacetic acid (PAA) to block viral DNA synthesis and late gene expression. Immunoblot at 72 hpi confirmed that UV irradiation blocked expression of all viral proteins tested, including IE1/IE2, the early/late protein pUL44, and the true late glycoprotein B (gB). In contrast, infection in the presence of either GCV or PAA had no effect on IE1, while IE2 and pUL44 levels were reduced, and, as predicted for a late protein, gB expression was fully blocked (Fig 3A). WDR5 protein and mRNA levels were compared during HCMV infection with or without GCV or PAA treatment. Compared to the mock-infected group, both WDR5 protein (Fig 3A) and mRNA levels (Fig. 3B) were elevated in virus-infected cells in both presence and absence of GCV or PAA. In HCMV-infected cells WDR5 protein levels were not significantly altered by GCV or PAA treatment.

**Fig 3.**
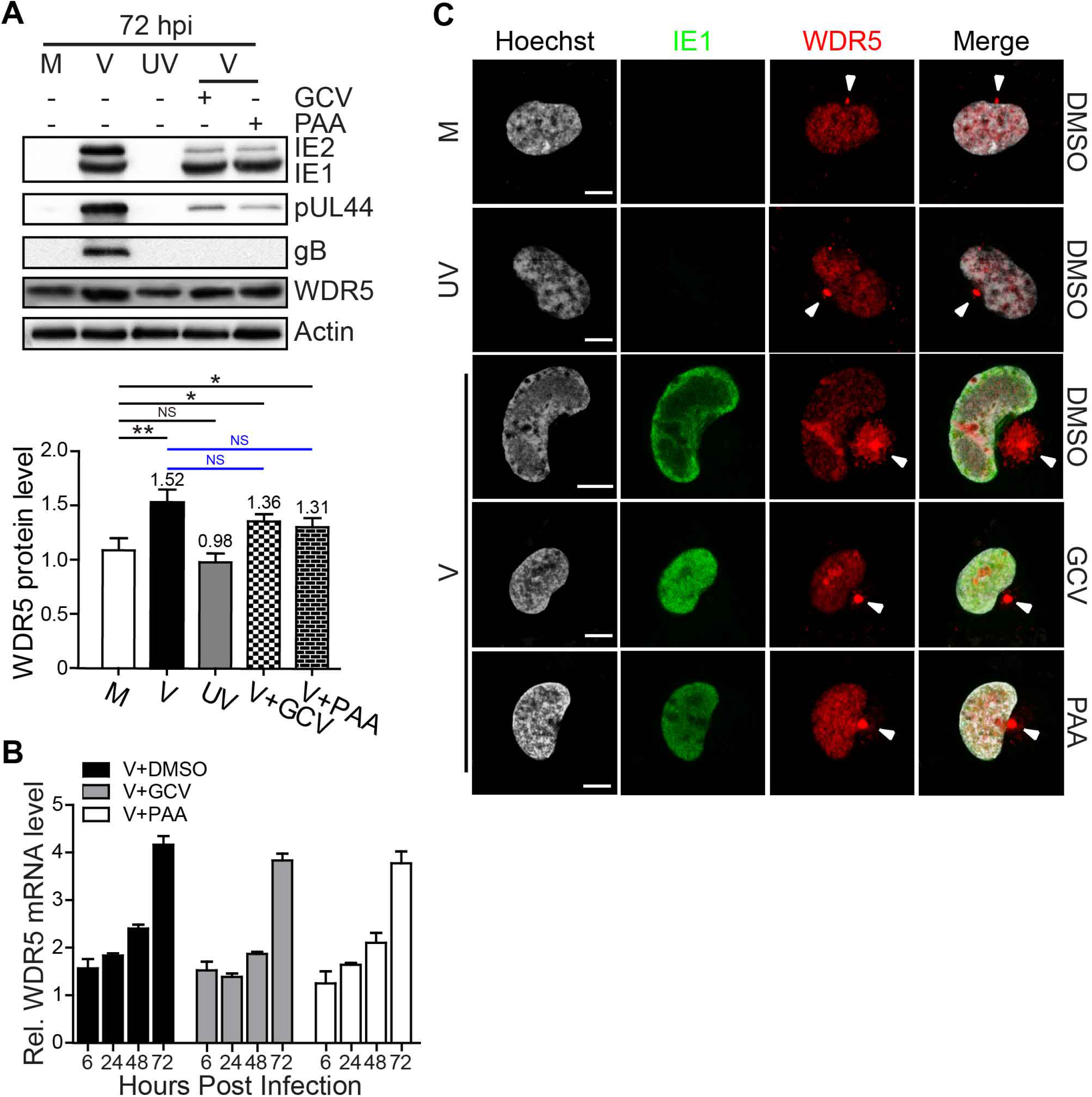
WDR5 perinuclear accumulation requires viral replication. HELFs were mock- (M), UV-inactivated HCMV (UV), or live HCMV-infected (V) at an MOI of 3 in the presence of DMSO, GCV (100 μg/ml), or PAA (100 μg/ml). At 72 hpi, samples were collected and analyzed. (A) The upper panel shows protein levels of WDR5 and viral proteins determined by IB with actin as a loading control. The lower panel shows the relative levels of WDR5 protein in infected vs. mock-infected cultures normalized with GAPDH. Data from three independent experiments were analyzed by one-way ANOVA, and the results are presented as averages +/− SD. *, P < 0.05; **, P < 0.01, NS, no significance. (B) Relative mRNA levels of WDR5. Samples were harvested at the indicated times, and total RNA was isolated. Relative mRNA levels of WDR5 were determined by qRT-PCR; GAPDH is an internal control. Data were normalized to levels in mock-infected cells to provide fold-changes post HCMV infection. (C) Accumulation of WDR5 in perinuclear region. WDR5 perinuclear foci are indicated by white arrows; scale bar = 10 μm.

We further investigated whether WDR5 perinuclear accumulation is altered by treatment of GCV or PAA. Characterization of replicate cultures by immunofluorescence revealed that WDR5 localization was similar to that in mock infected cells when virions were inactivated by exposure to UV. Both GCV and PAA dramatically inhibited perinuclear accumulation of WDR5 and perinuclear foci formation (Fig 3C). Although the expression of WDR5 was elevated by virus infection in the presence of GCV and PAA (Fig 3A and 3B), the perinuclear-accumulated WDR5 was still dramatically inhibited. These results indicated that WDR5 perinuclear accumulation and perinuclear foci formation are not due to increasing WDR5 levels, but are due to relocalization of WDR5 by a process that requires HCMV DNA synthesis and/or *de novo* synthesis of viral late proteins.

### Accumulation of WDR5 in perinuclear vACs requires microtubule polymerization

The location, morphology, and dependence on viral late gene expression suggested that WDR5 may be accumulating in the vAC. To confirm this possibility, colocalization of WDR5 with the vAC markers was examined. At 72 hpi, WDR5 accumulated in a juxtanuclear region in cells with kidney-shaped nuclei that also displayed cytoskeletal filaments (alpha-tubulin) radiating out to the rim of the plasma membrane (Fig 4A, a to d). This region also co-localized with gamma-tubulin (Fig 4A, e to h), which is the marker of the microtubule organizing center (MTOC) (48). This juxtanuclear region was wrapped around by a tightly nested ring of Golgi elements (anti-58K, Fig 4A, i to l) and colocalized well with early endosome marker EEA1 (Fig 4A, m to p). These characteristics have all been reported for the vAC (18), and confirmed that WDR5 accumulates in the vAC.

**Fig 4.**
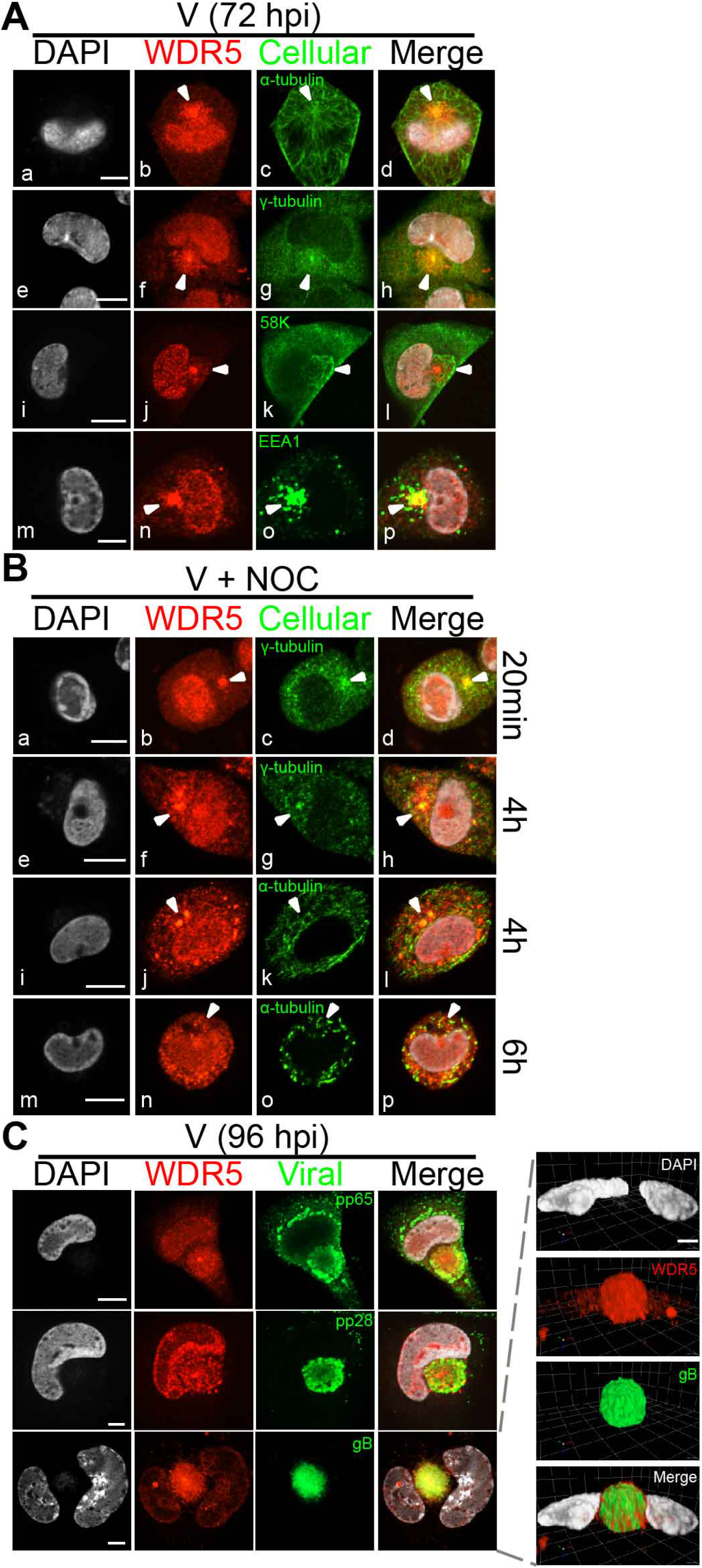
WDR5 localizes to vAC and MTOC. (A) HELFs were infected with HCMV at an MOI of 3 and at 72 hpi co-stained for WDR5 and markers of cytoskeleton (α-tubulin), centrosome (γ-tubulin), Golgi (58K), or early endosome (EEA1). (B) HELFs were infected for 72 h and then treated with 2 μM NOC for the times indicated prior to staining for WDR5. (C) HCMV-infected HELFs were stained at 96 hpi for WDR5 (red) and co-stained for viral proteins pp28, pp65, or gB (green). Panel below shows a 3D-reconstruction of WDR5 colocalization with gB from the cell shown in the panel above. Scale bar = 10 μm. (D) WDR5 associated with HCMV capsid protein in vAC. HFFs were infected with HCMV at an MOI of 3 and stained for WDR5 (green), SCP (red), and DNA (blue) at 96 hpi and analyzed with Nikon A1R HD25 confocal system. Top panels show deconvolved single channel maximum intensity z axis projections of a representative cell. Panels below show 3-D reconstructions of deconvolved confocal Z-series images of the vAC. All antibodies used in IFA were mouse monoclonal antibodies, nuclei were counterstained with DAPI. and scale bar = 10 μm.

To determine if WDR5 accumulation in the vAC was microtubule dependent similar to virion structural proteins such as pp150 (29), replicate cultures to those shown in Fig 4A were treated with nocodazole (NOC) to disrupt microtubule polymerization (49) and analyzed by IFA after 20 min, 4h, or 6h of NOC treatment. While γ-tubulin was less sensitive to NOC treatment (Fig 4B, c and g), α-tubulin filaments became fragmented and punctate in distribution after 4-6 h NOC treatment (Fig 4B, k and o). Concomitant with the fragmentation of microtubules, WDR5 perinuclear foci dispersed (Fig 4B, f, j and n).

In HCMV-infected cells, the vAC is enriched in various viral capsid, tegument, and envelope proteins (17, 18). To further confirm that accumulation of WDR5 in perinuclear foci resulted from recruitment of WDR5 to the vAC, infected cells were co-stained for WDR5 and viral vAC markers, including the tegument proteins pp28 and pp65 and the envelope glycoprotein gB (18). As shown in Fig 4C, pp28, pp65, and gB closely colocalized with WDR5. A three-dimensional (3-D) reconstruction from Z-stack images of the cell shown in the bottom panel of Fig 4C clearly demonstrated the colocalization of WDR5 with vAC marker gB. All these data indicated that WDR5 was driven to vAC during HCMV infection.

### WDR5 plays a crucial role in vAC formation and virion morphogenesis

To further explore whether WDR5 contributes to formation of vAC, we used a previously described pair of HELF-derived cell lines, one in which WDR5 is knocked down by shRNA (KD) and one expressing an irrelevant control shRNA (Ctl) (47). In KD cells, WDR5 knockdown altered formation and/or morphology of vAC, as determined by staining for gB. Irregular vAC exhibited multiple juxtanuclear foci, while in some infected cells no distinct vAC was observed (Fig 5A). Since proper vAC formation is necessary for efficient HCMV virion assembly and maturation. We further examine the virion morphogenesis by TEM in Ctl and KD cells. We previously have reported that knockdown of WDR5 dramatically decreased the number of matured virions in the vAC (47). As shown in Fig 5B (f to k), knockdown of WDR5 resulted in decrease of mature virions and distinct defects in virion maturation (white arrow). Nearly 39% of viral particles lacked envelope in KD cells (Fig. 5C). Inhibition of viral replication (Fig 3A and C) and disrupting microtubules (Fig 4B) both affect WDR5 accumulation in vAC, knock down WDR5 by shRNA altered formation and/or morphology of vAC (Fig 4D). All these data strongly support that WDR5 plays a crucial role in vAC formation and virion morphogenesis.

**Fig 5.**
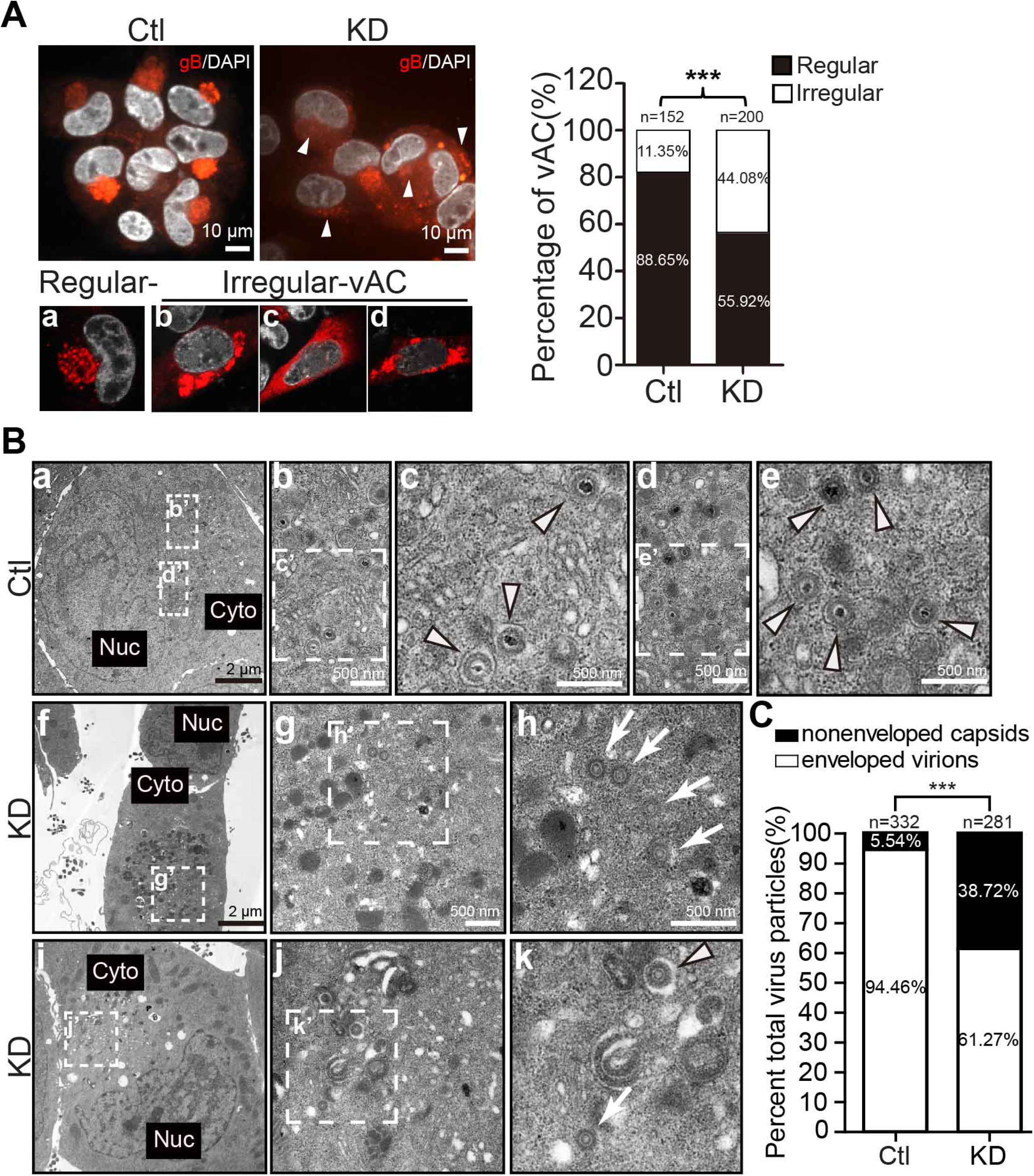
Knockdown WDR5 altered vAC formation and virion morphogenesis. (A) HELFs were transduced with shRNA to WDR5 (KD) or a scrambled control shRNA (Ctl) as described previously (47). Cells were infected with HCMV at an MOI of 0.5, harvested at 96 hpi, stained for gB with mouse monoclonal antibody, nuclei were counterstained with DAPI, scale bar = 10 μm (Left upper panels). Based on morphology, vACs were classified as regular or irregular, and representative images are shown (a, b, c, and d, Left lower panels). Regular and irregular vACs were quantitated (Right panel). n, number of quantitated infected cells; data were analyzed by chi-squared test. ***, P < 0.001.(B) Ctl (a to e) and KD (f to k) cells were infected with HCMV at an MOI of 0.5. At 120hpi, cells were fixed, embedded, sectioned and negative-stained for TEM analysis. Representative images are shown to illustrate the enveloped virions (arrowheads), and nonenveloped capsids (arrows) in the cytoplasm. (C) Quantitation and classification of viral particles. About three hundred total cytoplasmic viral particels were counted. The percentages of normal virions and nonenveloped capsids in each infection are illustrated. Data were analyzed by chi-squared test. n, number of cytoplasmic viral particles counted cells; ***, P < 0.001.

### The N-terminal domain of WDR5 is critical for WDR5 accumulation in vAC

The functional implementation of a protein depends on its subcellular distribution. To dissect which domain of WDR5 is critical for its accumulation in the vAC, a serial mutant of WDR5 were constructed based on its structure. WDR5 contains seven typical WD40 repeat domains (WD1 to WD7, Fig 6A), each WD40 repeat motif forms a blade β-propeller (37). A series of WDR5 deletion mutants were constructed as illustrated in Fig 6A. The distribution of Flag-tagged full-length WDR5 and the indicated mutants were determined. All the WDR5 constructs were transfected into HELF cells for 24 h, followed by either mock- or HCMV-infection at an MOI of 3. The full-length WDR5 showed a predominant nuclear localization in mock-infected HELFs, which is consistent with the results for endogenous WDR5 described above, accumulated in vAC in infected cells. Two mutants, Flag-aa1-82 and Flag-aa1-42, which retained the N-terminal amino acids 1-42, showed pan-cellular distribution upon mock-infection but still localized to vAC upon HCMV infection. In contrast, mutants lacking the amino acids 1-42 displayed mainly nuclear localization (*e.g.*, Flag-aa43-334) or pan-cellular distribution, and were not relocalized to vAC during HCMV infection. Taken together, these data indicate that the N-terminal amino acids 1-42 of WDR5 are critical for WDR5 accumulation in vACs.

**Fig 6.**
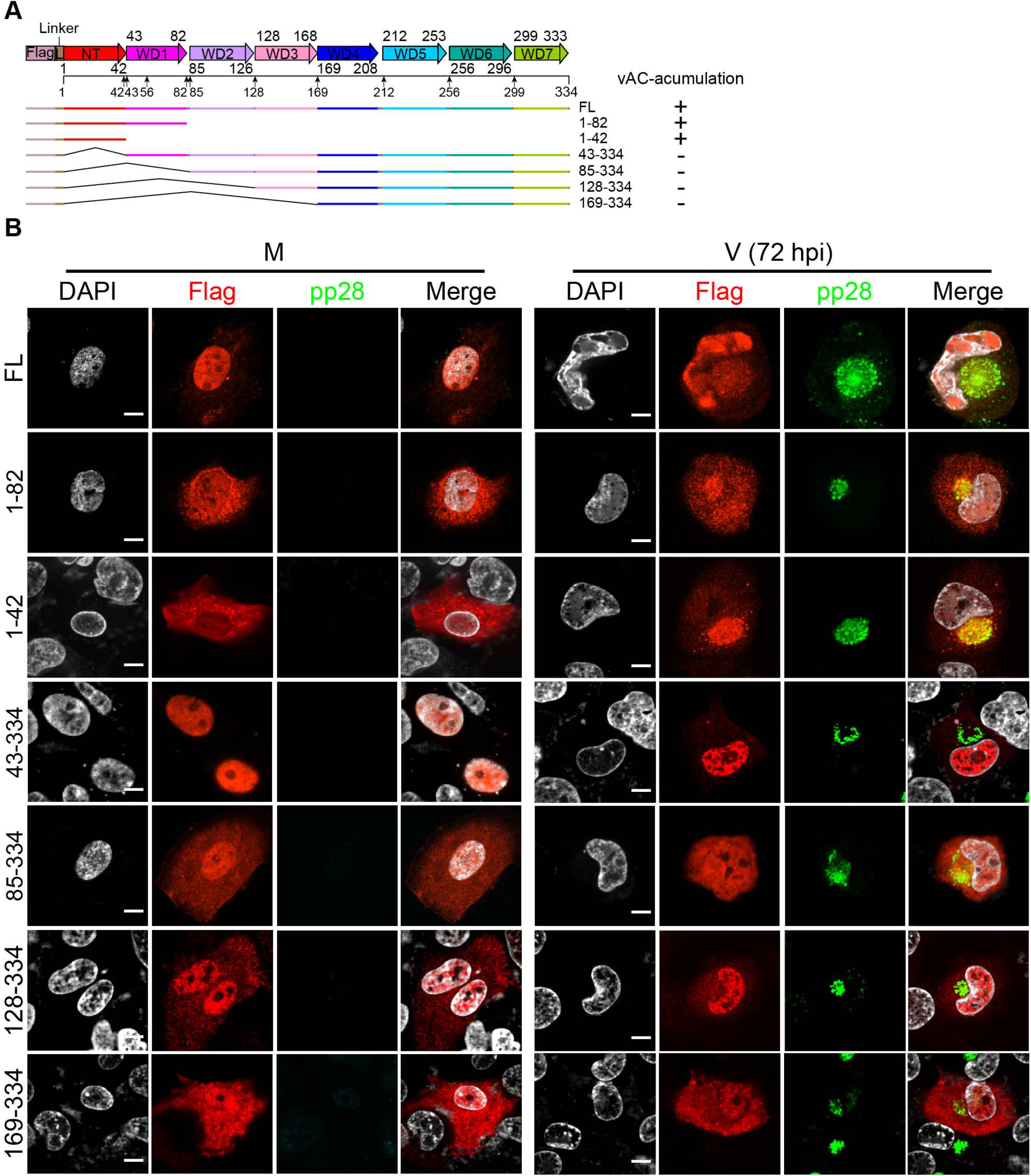
Subcellular localization of different WDR5 deletion mutants. (A) Schematic representation of N-terminal Flag-tagged full-length WDR5 and deletion mutants. Flag, Flag tag; L, linker-GAAAAS; NT, N-terminal; WD, WD40-repeat motif. (B) Expression vectors encoding full-length WDR5 or deletion mutants were transfected in HELF cells. At 24 hours post transfection, cells were mock-infected (M) or infected with HCMV (V) at an MOI of 3. At 72 hpi, samples were collected and analyzed by IFA for pp28 (green), Flag (red), or nuclei (DAPI, white). All antibodies used in IFA were mouse monoclonal antibodies. Scale bar = 10 μm.

### WDR5 co-fractionates with HCMV particles

Since WDR5 accumulated in the vAC, we next sought to determine if WDR5 is incorporated into viral particles. HELFs were mock- or HCMV-infected and cells were lysed by sonication. Intra- and extracellular particles were combined and then analyzed by iodixanol density gradient centrifugation, as illustrated in Fig 7A. Fractions were collected from top to bottom of the gradient, and analyzed by immuno-dot-blot for the presence of WDR5, gB, or pUL48, and for viral DNA by qPCR. WDR5 was undetectable in all fractions derived from mock-infected cultures (Fig 7B). In contrast, WDR5 was detected in fractions derived from HCMV-infected cultures and fractions with peak levels of WDR5 corresponded with those containing peak levels of gB, pUL48, and viral DNA (Fig 7B and 7C), suggesting that WDR5 is associated with HCMV particles.

**Fig 7.**
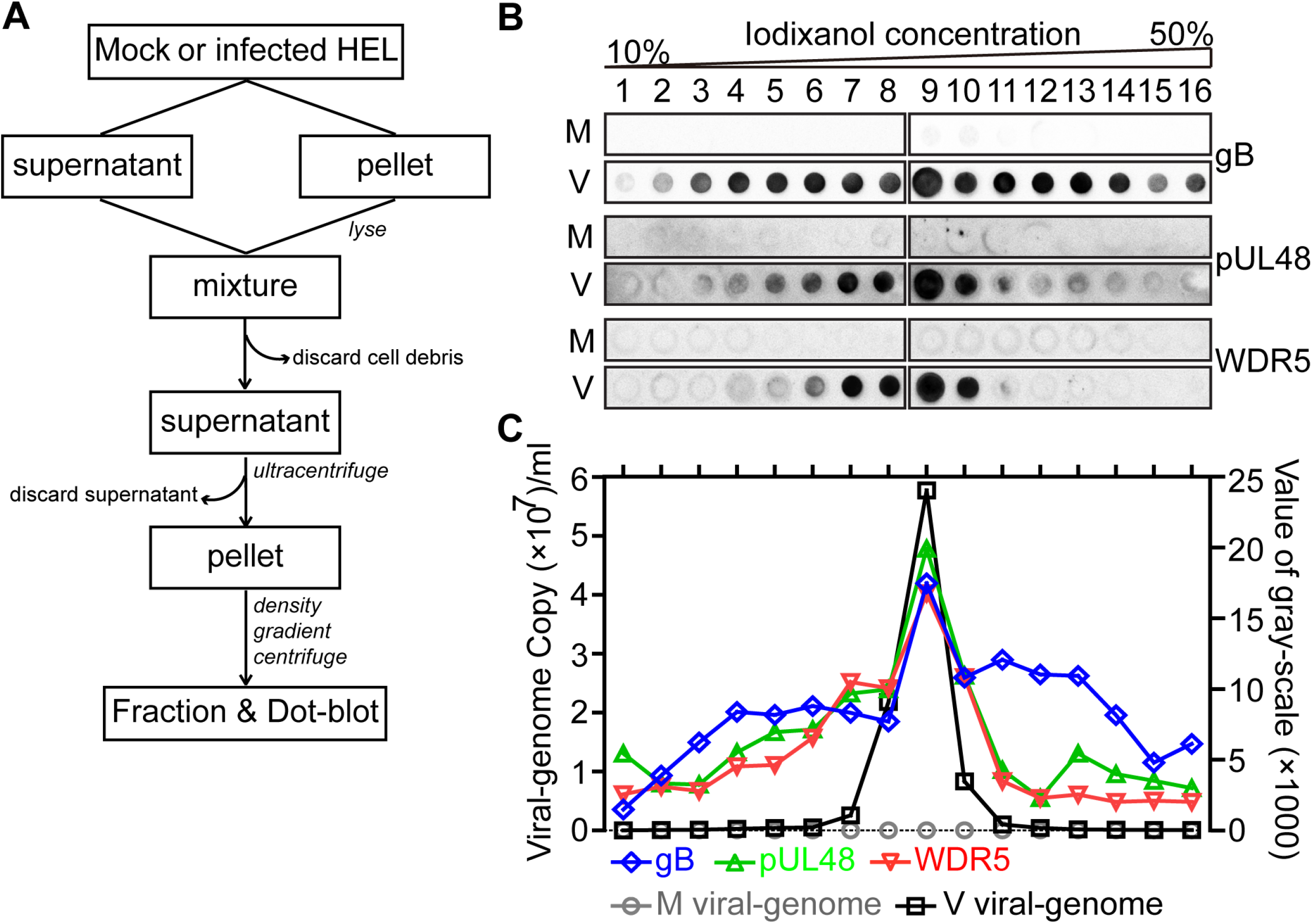
WDR5 co-purifies with HCMV particles by gradient fractionation. (A) Flow chart illustrating the analysis of particles from mock- or HCMV-infected cultures by iodixanol density gradient centrifugation (see Materials and Methods for details). (B) Gradient fractions derived from mock-infected (M) or infected (V) cultures were analyzed by immune-dot-blot for WDR5, gB, or pUL48. (C) Viral-genome copies in each fraction were determined by qPCR and plotted together with protein levels quantitated using ImageJ software from the dot-blots shown in (B).

### WDR5 is present in the tegument layer of HCMV virions

To determine whether WDR5 is incorporated into virions, extracellular virions were purified from infected-cell culture supernatants as described previously (50). The purified virions were then either mock treated or treated with 2% Triton X-100 at room temperature for 1 h, layered onto a 10% to 50% iodixanol continuous gradient, and ultra-centrifuged at 100,000 × *g* for 2 h. Fractions were collected from top to bottom of the gradient and analyzed by IB for WDR5, structural viral proteins, including major capsid protein (MCP), tegument proteins pp28, pp65, pp71, pIRS1, pTRS1, pUL48, and the envelope glycoprotein B (gB), as well as the non-structural viral proteins IE1/2 and pUL44. IE1/2 and pUL44 were present in infected-cell lysates but were not in any gradient fractions, confirming effective separation of virions from infected-cells (Fig 8A). In gradient fractions of mock-treated virions, fractions 9 and 10 contained peak levels of MCP, tegument proteins, gB, and WDR5 (Fig 8A, upper panel). In gradient fractions of Triton X-100-treated virions, peak levels of MCP and tegument proteins shifted to fraction 11, while the majority of gB and a fraction of the outer tegument protein pp28 shifted to lower density fractions and no longer co-fractionated with MCP (Fig 8A, lower panel). These results suggested that Triton X-100 treatment removed the envelope and some of the outer tegument layers from the virion particles. The finding that WDR5 remained predominantly in fraction 11 following Triton X-100 treatment (Fig 8A, lower panel) suggested that WDR5 was either associated with the inner tegument layers or was directly attached to capsids.

**Fig 8.**
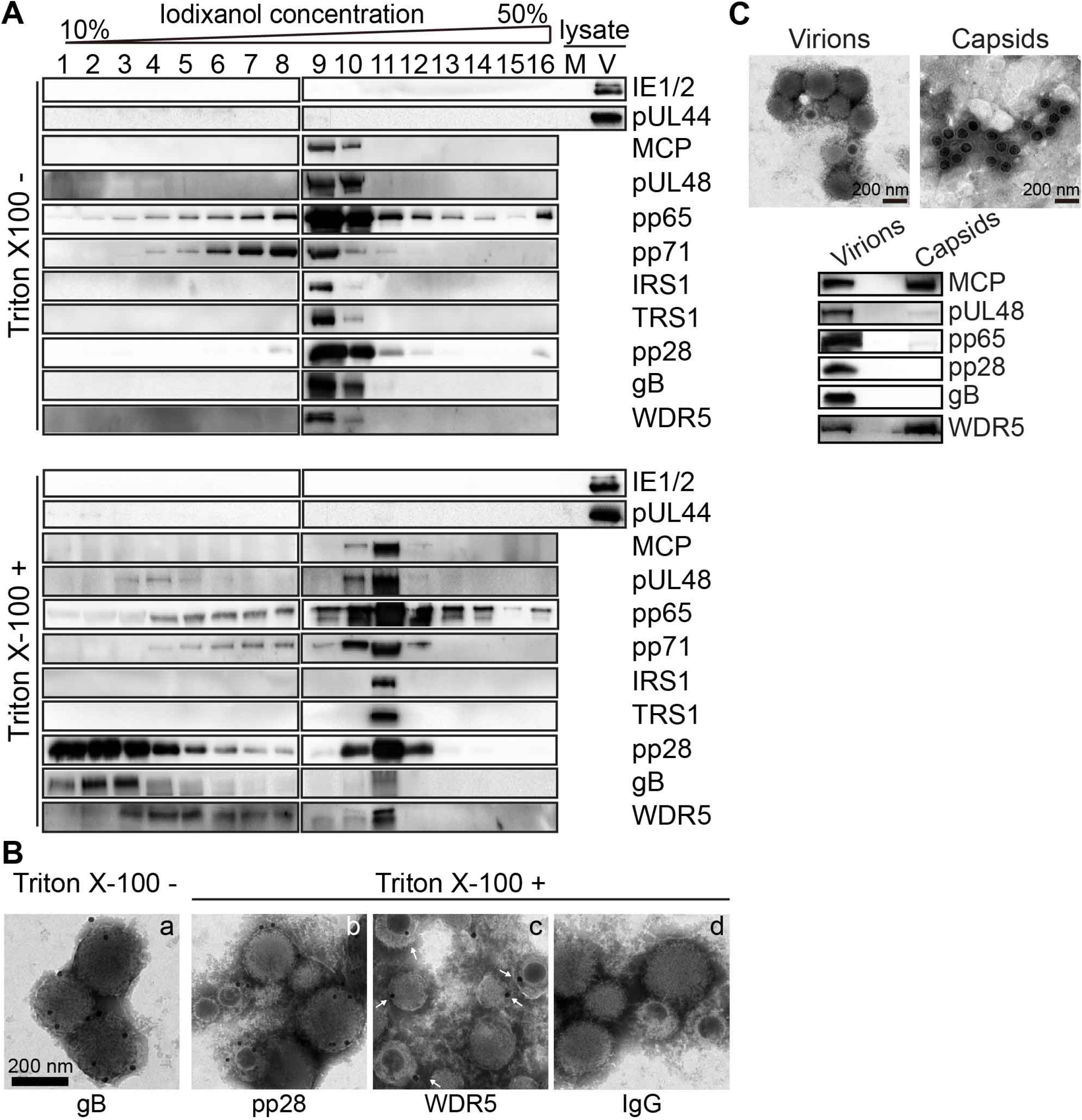
WDR5 is present in HCMV virions. (A) Purified HCMV virions were mock treated or treated with 2% Triton X-100 for 1 hour at room temperature then analyzed by iodixanol density gradient centrifugation. Fractions collected from gradients of virions that were mock treated (top) or Triton X-100-treated (bottom), as well as unfractionated lysates from mock- (M) or HCMV-infected (V) HELFs, were analyzed by IB using the indicated antibodies. (B) Mock- or Triton X-100-treated virions were analyzed by immuno-gold electron microscopy using mAbs to gB (a), pp28 (b), WDR5 (c), or normal mouse IgG (d). Samples were negatively stained with phosphotungstic acid and visualized by electron microscopy (25,000 × magnification). (C) HCMV virions were purified as previously described. Nuclei from infected-HELs were prepared at 60 hpi and intracunclear capsids were purified as in described in Materials and Methods. Purified virions and capsids were visualized by electron microscopy. The components (viral proteins and WDR5) were analyzed by IB.

Immunogold labeling was next used to further examine WDR5 association with virions. Purified virions were again either mock treated or treated with Triton X-100 (as described above) to allow access of antibodies to proteins in the inner layers of treated viral particles. The particles were then labeled with mAbs to gB, pp28, or WDR5 followed by immunogold labeling and detection by electron microscopy. Consistent with the established presence of gB in the virion envelope, gold particles were observed decorating the surface of mock-treated particles labeled using anti-gB antibody (Fig 8B, panel a). In contrast, and consistent with the known localization of pp28 in the tegument of the virion, gold particles were observed decorating the surface of Triton X-100-treated particles labeled using anti-pp28 antibody (Fig 8B, panel b). Immunogold labeling with anti-WDR5 antibody was similar to that of pp28, suggesting that WDR5 was localized within the tegument layer (Fig 8B, panel c). Moreover, capsids purified from infected-cell nuclei (before nuclear egress) or from virions purified from culture supernatants both contained WDR5 (Fig. 8C).

Taken together, these results indicate that WDR5 is associated with capsids before nuclear egress, translocates to and accumulates in vAC, finally incorporates into virion tegument layer, which is critical for virion morphogenesis.

### WDR5 interacts with viral structural proteins

The above findings indicate that WDR5 is associated with capsids before nuclear egress, then accumulates in the vAC, and is ultimately incorporated into virions. We therefore infer that WDR5 likely interacts with one or more viral structural proteins. To test this hypothesis, Flag-tagged WDR5 was overexpressed in HELF cells followed by mock- or HCMV-infection for 96 h. Cell lysates were immunoprecipitated (IP) with anti-Flag antibody and subjected to SDS-PAGE and Coomassie staining. There were four different distinct protein species (Fig. 9A, red arrows) in the viral-infected immunoprecipitates when compared to the mock control. These four species were further analyzed by liquid chromatography-tandem mass spectrometry (LC-MS/MS) to identify and quantify the proteins. As shown in Fig 9B, viral structural proteins, including MCP, pp150, pp65, pIRS1, and pTRS1, were identified in the immunoprecitates.

**Fig 9.**
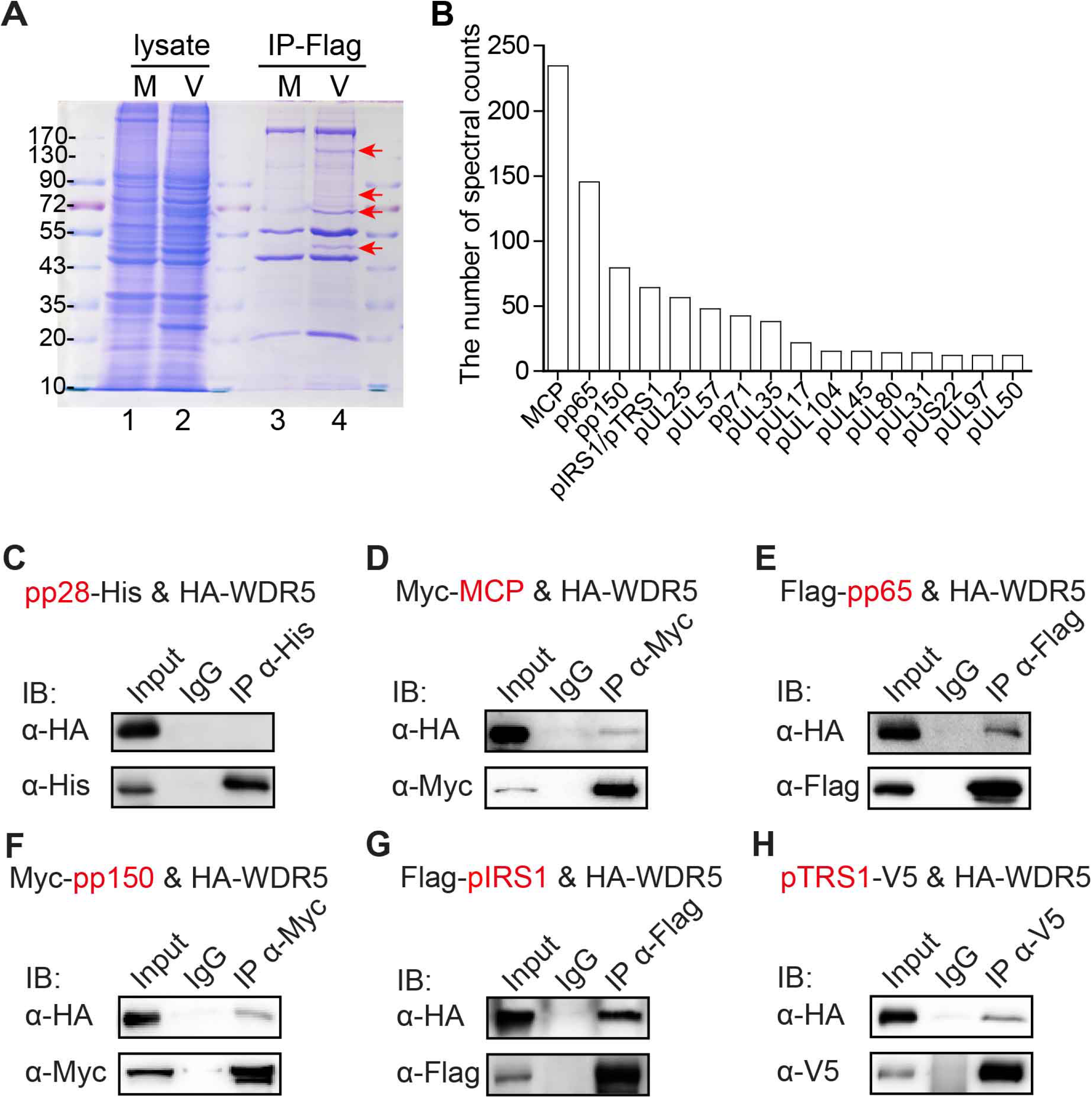
WDR5 interacts with MCP, pp150, pp65, pIRS1, and pTRS1. (A) HELF overexpressing with Flag-tagged WDR5 were mock- (M) or HCMV-infected (V) at an MOI of 3. At 96 hpi lysates were immunoprecipitated by anti-Flag antibody, separated by SDS-PAGE, and stained with Coomassie Brilliant Blue. Four protein species specific to infected cells (red arrows, lane 4) were cut from the gel and combined and analyzed by LC-MS/MS. (B) Spectral counts of identified viral proteins by mass spectrometric analyses are shown. (C-H) HEK293T cells were co-transfected with a plasmid expressing HA-tagged WDR5 (HA-WDR5) along with plasmids expressing His-tagged pp28 (pp28-6×His) (C), Myc-tagged MCP (6×Myc-MCP) (D), Flag-tagged pp65 (Flag-pp65) (E), Myc-tagged pp150 (6×Myc-pp150) (F), Flag-tagged pIRS1 (Flag-IRS1) (G), or V5-tagged pTRS1 (pTRS1-V5) (H). After 48 h lysates were immunoprecipitated and then immunoblotted using the indicated antibodies.

To confirm WDR5-viral protein interactions, HEK293T cells were co-transfected with plasmids expressing HA-tagged WDR5 and each of the candidate viral protein interactors (Myc-MCP, Flag-pp65, Myc-pp150, Flag-pIRS1, or V5-pTRS1). Each viral protein was immunoprecipitated using their respective epitope tags and the immunoprecipitates were then probed for WDR5 by IB for the HA tag. WDR5-HA could be detected in IPs for MCP, pp65, pp150, pIRS1, and pTRS1 but not for pp28 or the IgG controls (Fig. 9C-H). Taken together, these results indicate that WDR5 interacts with viral structural proteins, which explains its accumulation in vAC and its role in facilitating mature virion morphogenesis.

## Discussion

During the course of a viral infection, cellular proteins can be incorporated into viral particles. For example, both LC-MS/MS and multidimensional protein identification technology (MudPIT) have been used to identify components of influenza virions, and while seventeen virion-associated cellular proteins were identified by both methods, six others were identified only with the LC-MS/MS analysis and 13 were identified only with MudPIT analysis (51), suggesting that multiple approaches may be necessary to identify all of the virion-associated cellular factors. In some cases, cellular proteins are incorporated randomly and have not been shown to play a role in the infectivity of the mature virion, whereas in other cases cellular proteins are selectively incorporated into virions through specific interactions with virion structural proteins or the viral genome. With regard to HCMV, LC-MS/MS has identified over 70 host proteins in virions. These cellular proteins function in diverse processes, including ATP and Ca^2+^ binding, chaperones, cytoskeleton, enzymatic activities, glycolysis, protein transport, signal transduction, transcription, and translation (31). Some have been reported to function during certain stages of the infection process, suggesting that these proteins are incorporated into the virion to facilitate virus infectivity and replication (31). Others may serve to evade host immune mechanisms; for example, in the late stage of infection intracellular viral DNA sensors and restriction factors such as IFI16 traffic to vAC where they are incorporated into mature virions (52).

Previously, we found that HCMV infection increases WDR5, and WDR5 facilitates capsid nuclear egress (47). In the present study we analyzed the cellular distribution of WDR5 and observed that the distribution of WDR5 was altered by HCMV infection, as it accumulated in vAC. Treatment with viral replication inhibitors (GCV or PAA) and microtubule polymerization inhibitor (NOC) significantly prevented WDR5 accumulation in the vAC. Inhibition of viral replication, including viral DNA replication and expression of late viral structure proteins, impairs formation of the vAC. The vAC formation is also dissolved by microtubule polymerization inhibitor. Under both conditions, the protein levels of WDR5 are not significantly changed, but vAC formation is dramatically impaired. Thus, relocation and accumulation of WDR5 in vAC requires HCMV late gene expression and depends on microtubule polymerization. Taken together, these findings suggest that WDR5 accumulation in the vAC occurs as a result of a specific HCMV-dependent mechanism, although this mechanism remains undefined.

In order to determine if WDR5 was incorporated into virions, we analyzed intra- and extracellular viral particles and purified virions for the presence of WDR5 after purification of virion particles by density gradient centrifugation. In each case WDR5 cofractionated with markers for virions or capsid particles but was not detected in gradient fractions derived from uninfected cultures. Furthermore, we showed that WDR5 interacts with virion tegument proteins pp65, pIRS1, and pTRS1. Therefore, we propose that WDR5 resides within the tegument layer of HCMV virions. To investigate the location of WDR5 in virions, we treated virions with Triton X-100, a detergent that solubilizes virion membranes. Following Triton X-100 treatment, a significant amount of WDR5 remained associated with the detergent treated virions. In addition, immuno-gold electron microscopy indicated that WDR5 was present within virions and was localized to the tegument. These data, together with our previous findings that WDR5 plays a role in the assembly of the NEC and capsid egress, suggest that during nuclear egress WDR5 may initially enter the perinuclear cisterna together with capsids and travels along with capsids to the cytoplasm and vAC (Fig 10). WDR5 then interacts with viral structural proteins (*e.g.,* MCP, pp150, and pp65), thereby accumulating in vAC; finally, WDR5 is packaged into virions along with numerous tegument proteins and glycoproteins.

**Fig 10.**
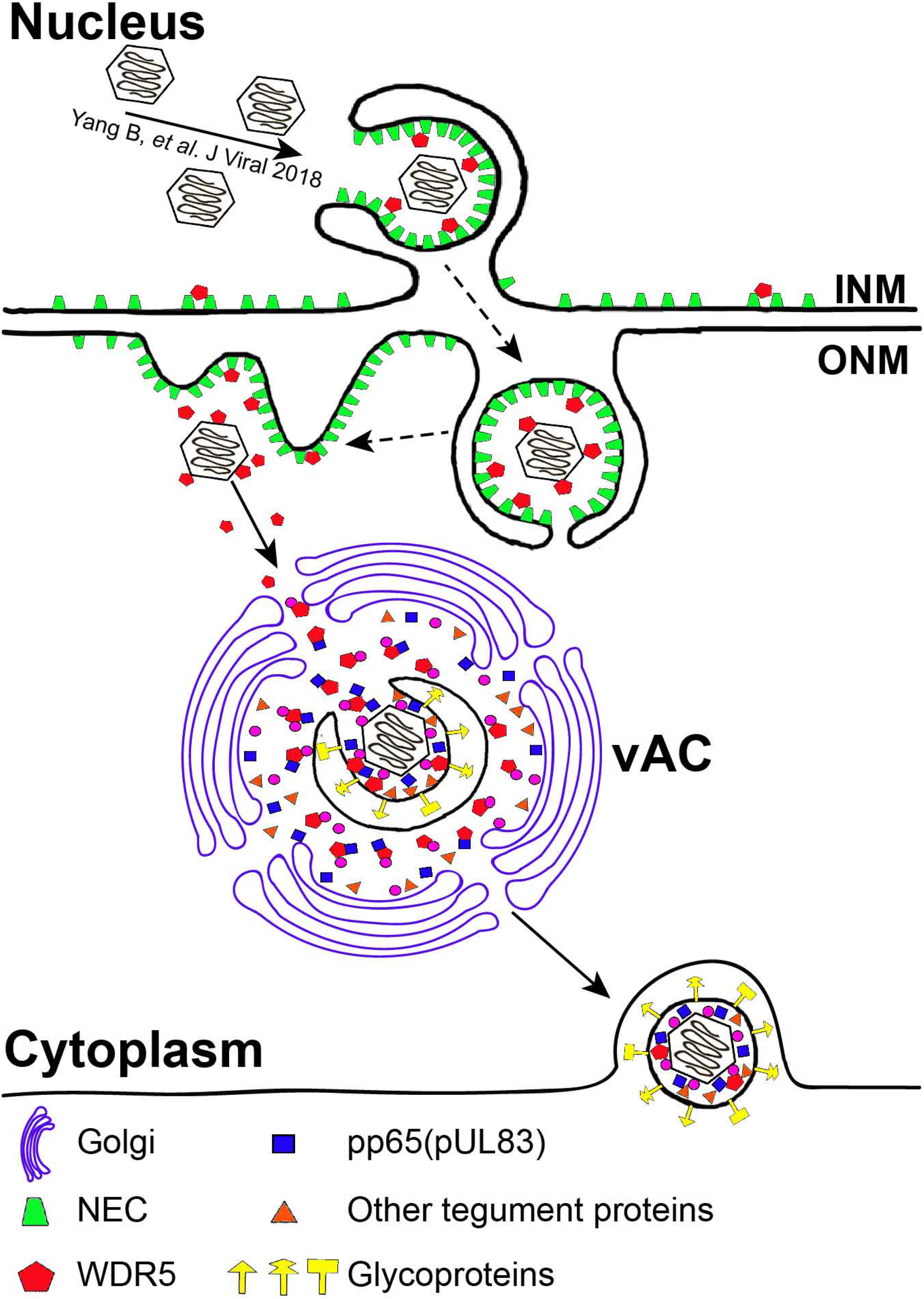
Working model. In the nucleus WDR5 facilitates capsid nuclear egress by promoting assembly of the NEC (Yang B, *et al*. J Virol 2018). WDR5 may egress from the nuclei with the capsids and then be transported to the vAC together with capsids and/or tegument proteins (*eg.* pp150 and pp65, *etc*.). In the vAC, WDR5 is incorporated into maturing virions.

In summary, the data provided in this study suggest a dual role for WDR5 during HCMV assembly. During nuclear egress, it modulates NEC assembly and facilitates capsid nuclear egress. Subsequently, it translocates along with capsid from the nucleus into the cytoplasm by interacting with MCP; it accumulates in vAC by interacting with multiple late viral structures proteins and is incorporated into virions. Since WDR5 interacts with multiple viral structure proteins, it could function during virion morphogenesis by stabilizing multi-protein complexes. WDR5 plays a critical role in capsid nuclear egress, vAC formation, and virion assembly. Our findings further demonstrate that HCMV manipulates existing cellular mechanisms, including the localization of cellular proteins, to facilitate its replication, assembly, and egress. These data highlight that WDR5 is a potential new target for antiviral drug development.

## Acknowledgements

We thank Dr. Hua Zhu from Rutgers-New Jersey Medical School for providing VZV Oka strain, Qiyi Tang from Howard University College of Medicine for providing MCMV K181 strain, and Drs. Zhengli Shi and Zhihong Hu from Wuhan Institute of Virology, CAS, for providing GFP-expressing AdV and AcMNPV, respectively. We thank Dr. Wade Gibson from Johns Hopkins University for providing pUL48 rabbit serum, Dr. Yan Zhou from Wuhan University for providing γ-Tubulin antibody, and Dr. Thomas E. Shenk from Princeton University for providing pIRS1 and pTRS1 antibodies. We thank Dr. Adam P. Geballe from Fred Hutchinson Cancer Research Center, University of Washington for providing pTRS1 expressing plasmid. We thank Ding Gao, Anna Du, and Pei Zhang of The Core Facility, Wuhan Institute of Virology, CAS, for technical support of electron microscopy.

## Materials and Methods

### Ethics statement

Human embryonic lung fibroblast cells (HELs) were isolated from postmortem embryo lung tissue. The original source of the anonymized tissues was Zhongnan Hospital of Wuhan University (China). The cell isolation procedures and research plans were approved by the Institutional Review Board (IRB) (WIVH10201202) according to the Guidelines for Biomedical Research Involving Human Subjects at Wuhan Institute of Virology, Chinese Academy of Sciences. The need for written or oral consents was waived by the IRB (53).

### Cells and cell culture

HELs were isolated and maintained as described previously (53). HELF, kindly provided by Dr. Jason J. Chen at Columbia University, are human embryonic lung fibroblasts that have been retrovirally transduced with human telomerase (hTERT). Clonal cell lines isolated from HELFs that were transduced with WDR5 shRNA (KD) and scrambled shRNA (Ctl) were described previously (47). Both HELs and HELF were cultured in Minimum Essential Medium (MEM, Cat.#41500-034, Thermo Fisher) supplemented with 10% fetal bovine serum (FBS, Cat.#10099-141, Thermo Fisher) and penicillin-streptomycin (100 U/ml and 100 μg/ml, respectively; Cat.#15140-122, Thermo Fisher). Human embryonic kidney 293T cells (HEK293T) were purchased from ATCC (CRL-11268) and cultured in Dulbecco’s Modified Eagle Medium (DMEM; Cat.#11995-123, Thermo Fisher) supplemented with 10% FBS and penicillin-streptomycin, as above. Cells were cultured at 37 °C in a humidified atmosphere containing 5% CO_2_.

### Viruses

HCMV Towne strain (ATCC-VR 977) was used in this study. HSV-1 strain H129 expressing GFP was previously constructed by our laboratory (54). VZV Oka strain expressing GFP (55) was a gift from Dr. Hua Zhu, Rutgers-New Jersey Medical School, and MCMV strain K181 was a gift from Qiyi Tang, Howard University College of Medicine, USA. GFP-expressing AdV and AcMNPV were provided by Dr. Zhengli Shi and Zhihong Hu, Wuhan Institute of Virology, Chinese Academy of Sciences, respectively.

### Plasmid construction

Primers used for plasmid construction are shown in Table 1. WDR5 cDNA (GenBank #NM_017588) was derived by reverse transcription as described previously (47). The cDNA was PCR amplified then cloned into either *Eco*RI/*Xho*I-restricted pCMV-HA to generate plasmid HA-WDR5 or into *Xba*I/*EcoR*I-restricted pCDH-Flag to generate Flag-WDR5. The *UL83* ORF encoding pp65 and *IRS1* ORF encoding pIRS1 were PCR amplified from Towne-BAC genome (56) and cloned into *Bam*HI/*Kpn*I-restricted pXJ40-Flag to generate plasmid Flag-pp65 and Flag-pIRS1, respectively. The *UL99* ORF encoding pp28 was PCR amplified and cloned into *Bam*HI/*Xba*I-restricted pcDNA3.1-V5/6×His to generate plasmid pp28-6×his. The *UL86* ORF encoding MCP and *UL32* ORF encoding pp150 were amplified from Towne-BAC genome (56) and inserted into a vector with 6×Myc by cloning recombination (ClonExpress II One Step Cloning Kit, Cat.#C112, Vazyme). The *TRS1* ORF encoding pTRS1 fused with a carboxyl-terminal 6×His epitope tag and V5 tag was a gift from Dr. Adam P. Geballe, Fred Hutchinson Cancer Research Center and University of Washington, which have been previously described as pEQ981 (57). All the primers used in this study are listed in Table 1.

**Table 1.**
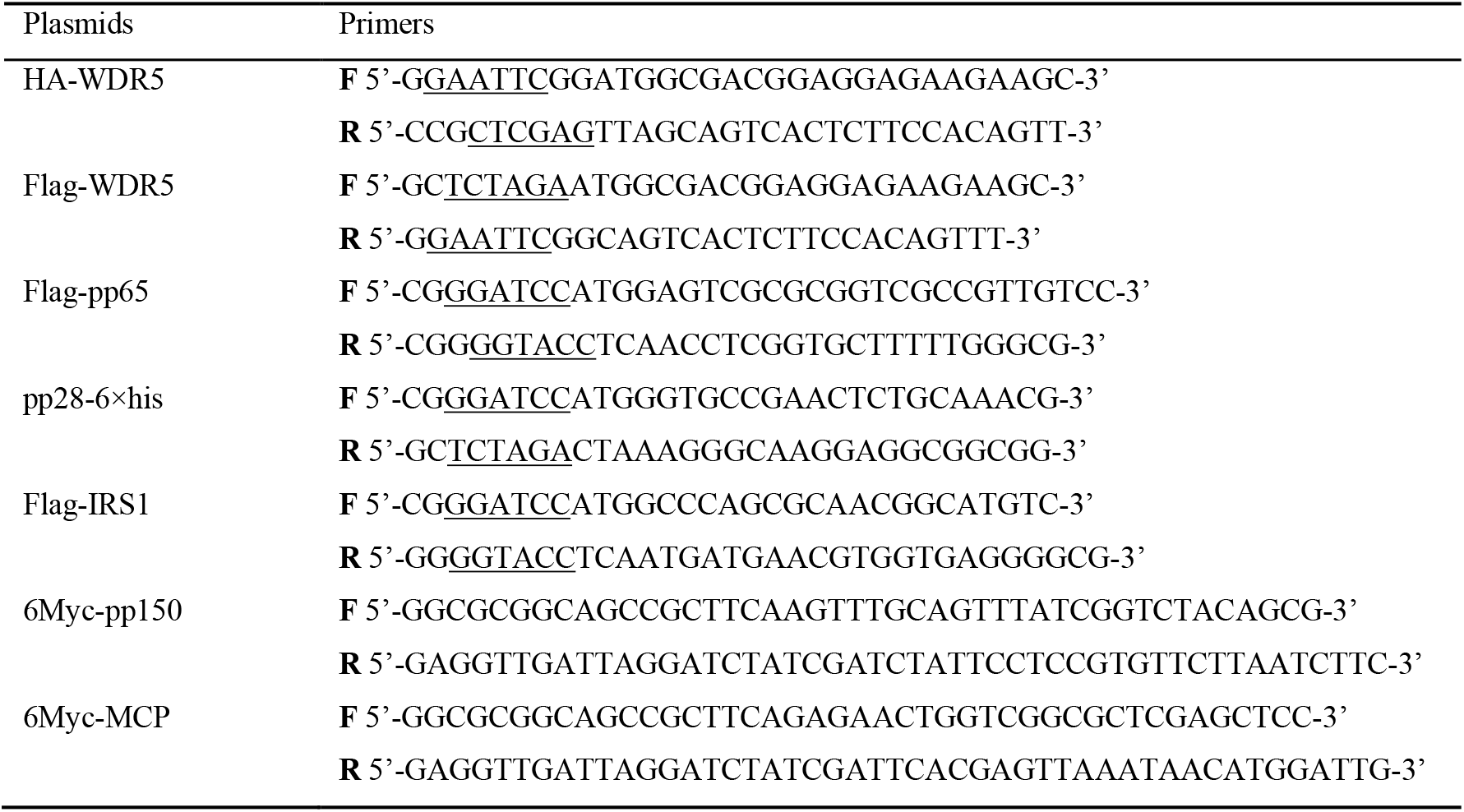
Primers used in this study.

### Transient transfection

1.5×10^6^ HEK293T cells were seeded into 100-mm dishes. The next day the medium was changed 2h prior to transfection via Ca_2_(PO_4_)_2_ precipitation with 10 μg HA-WDR5 along with 10 μg pp28-his, Myc-MCP, Flag-pp65, Myc-pp150, Flag-pIRS1, or pTRS1-V5, as described previously (53). 2×10^6^ HELFs were seeded onto coverslips in 12-well plates. Medium was changed at 2h prior to transfection in the next day. Plasmids (1.2 μg for each well) expressing full-length and truncated WDR5 were transfected into HELFs using Lipofectamine 2000 reagent (Cat.#11668-019; Thermo) according to the manufacturer’s instructions.

### Quantitative reverse transcriptase PCR (qRT-PCR)

HELFs were infected at an MOI of 3 and harvested at the indicated times post infection. A total of 1 × 10^6^ cells were used for total RNA extraction using RNAiso Plus reagent (Cat.#9109, TaKaRa), followed by treatment with 10 U of recombinant DNase I (Cat.#2270A, TaKaRa) to remove residual DNA. One μg RNA of each sample was reverse transcribed with a RevertAid H Minus first-strand cDNA synthesis kit (Cat.#K1631, Fermentas) with random primers. Then qPCR was performed on a real-time thermocycler (Bio-Rad; Connect) using SYBR green PCR master mix (Cat.#4309155, Applied Biosystems) in 20 μl reactions for 40 PCR cycles as described previously (58). The PCR primers for WDR5 were: 5’-GGTGGGAAGTGGATTGTGTC-3’ and 5’-GCAGCAGAGGCGATGATG-3’. The PCR primers for GAPDH were: 5’-GAGTCAACGGATTTGGTCGT-3’ and 5’-GACAAGCTTCCCGTTCTCAG-3’.

### IB

Cells were harvested in cell lysis buffer (Cat.#P0013, Beyotime) containing protease inhibitor cocktail (Cat.#04693159001, Roche) and homogenized by ultrasonication. Protein concentrations of lysates were determined by Bradford assay (Cat.#500-0205, Bio-Rad). After boiling with loading buffer, cell lysates containing equal protein amounts were separated by SDS-PAGE and transferred to PVDF membranes (Cat.#ISEQ00010, Millipore). Membranes were sequentially probed with primary antibodies and appropriate peroxidase-conjugated secondary antibodies. All antibodies used for IB are listed in Table 2. The viral antibodies for IE1 (59), pUL48 (60), pIRS1 (61), pTRS1 (61) and MCP (62) and mIE1 (63) were described previously. Blots were developed using the SuperSignal West Femto Chemiluminescent Substrate (Cat.#34095, Thermo Fisher), signals were detected using a FluorChem HD2 System (Alpha Innotech), and quantification was performed using ImageJ software (NIH).

**TABLE 2.**
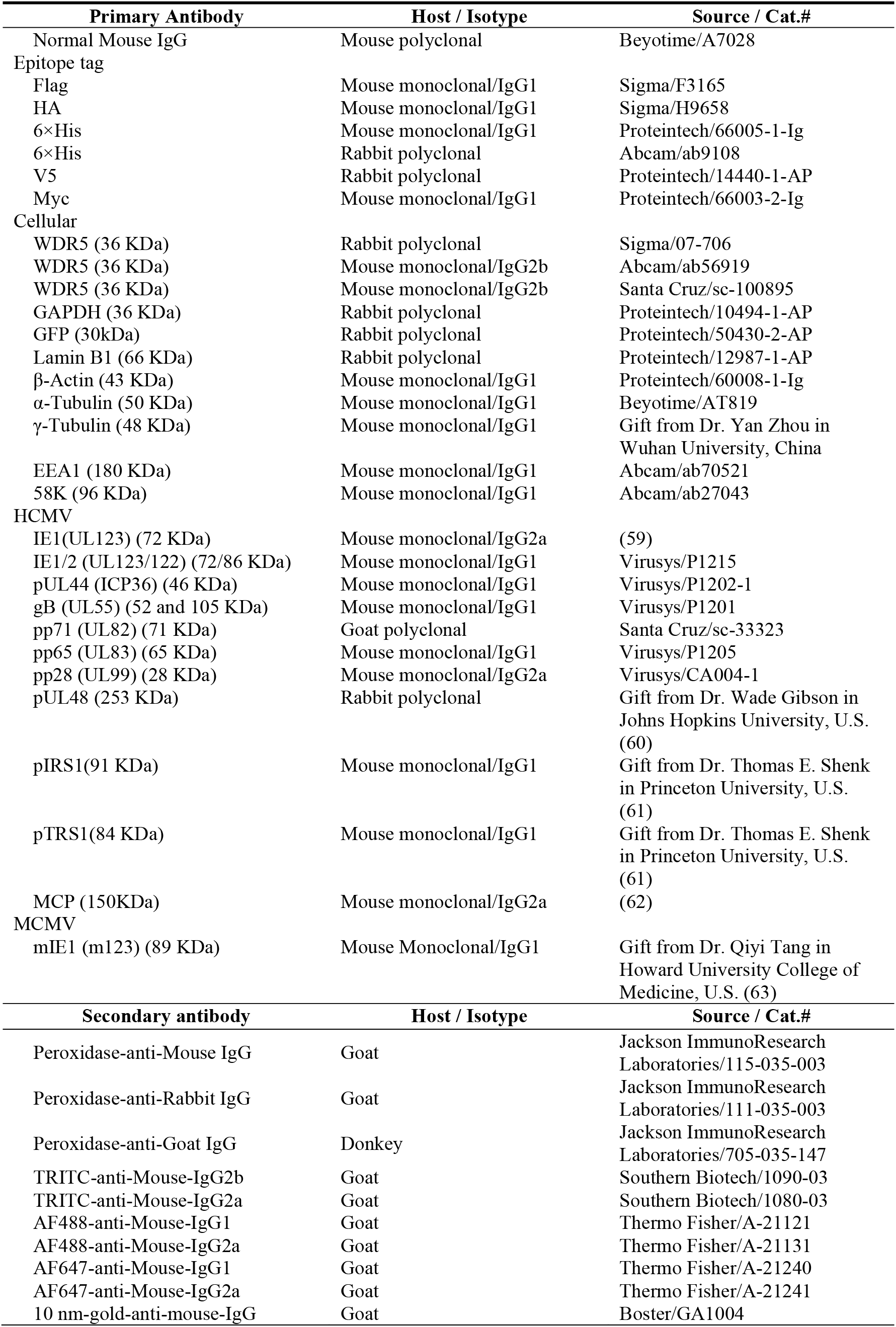
Primary and Secondary Antibodies Used in IFA and IB.

### IP

HEL cells were mock infected or infected with HCMV strain Towne at an MOI of 3 and harvested at 72 hpi. Cells were lysed in IP lysis buffer (Cat.#P0013, Beyotime) for 1h at 4 °C then centrifuged at 120,000 × g for 5 min to remove cell debris. Samples were mock treated or nuclease treated by incubation with 10 mM MgCl_2_, 100 μg/ ml Dnase I (Cat.#2270A, TaKaRa) and 100 μg/ml Rnase A (Cat.#AC118, OMEGA) for 1 h at 37 °C. IP was performed by incubation of the resulting lysates overnight at 4 °C with mouse monoclonal anti-Flag antibody (Cat.# F3165, Sigma) or with normal mouse IgG (Cat.#A7028, Beyotime). Protein A+G agarose beads (Cat.#P2012, Beyotime) were added and incubated for 3 h at 4 °C and then washed five times with lysis buffer. Loading buffer was added and samples were boiled for 5 min before separation by SDS-PAGE followed by IB for detection of WDR5, pp65, or pp28, as described above.

### IFA

HELFs or HFFs were seeded onto coverslips and after attachment infected with HCMV at an MOI of 3. Coverslips were collected at the indicated times post infection and fixed with 4% paraformaldehyde. Target proteins were detected by incubation with primary antibodies and appropriate secondary antibodies, as listed in Table 2, and as described previously (47). Nuclei were counterstained with DAPI (Cat.#D9542, Sigma). Images were obtained using a UltraVIEW VoX (Perkin Elmer) spinning disk laser confocal scanning microscope. 3D z-axis image stacks were acquired with 0.2-μm spacing and 3D modeling was performed using Volocity 5.5 software (Perkin Elmer). Images were median-filtered to reduce noise and contrast was enhanced to improve resolution.

### Cytosolic and nuclear fractionation

HELFs were mock infected or infected with HCMV at an MOI of 3, harvested at 72 hpi, and fractionated using the Qproteome Cell Compartment kit (Cat.#37502, Qiagen) as described previously (64). Fractions were analyzed by IB using GAPDH and Lamin B1 antibodies to confirm separation of cytosolic and nuclear fractions, respectively.

### Quantitation of viral genome copy number

Genomic DNA from each fraction were analyzed by qPCR to quantitate viral DNAs using HCMV UL83 primers as described previously (53). Means and standard deviations (SD) from at least three independent experiments were calculated.

### Virus particles and capsid purification

HELs were infected with HCMV at an MOI of 0.02 and harvested after CPE reached 100%. Culture medium was collected and clarified by centrifugation at 2,000 × *g* for 10 min. Virus particles were pelleted from the supernatants at 100,000 × *g* for 2 h at 4 °C in an SW 32Ti rotor using a Beckman Optima™ L-100 XP Ultracentrifuge. The pellet was resuspended and layered onto an iodixanol gradient that was prepared by sequentially layering 800 ul volumes of 50%, 40%, 30%, 20%, and 10% iodixanol (Cat.#D1556, Sigma) in PBS (v/v) followed by incubation overnight 4 °C. Gradients were then centrifuged 100,000 × *g* for 2 h at 4 °C by using an SW 55Ti rotor and Optima™ MAX-XP Ultracentrifuge. The virion-containing band was observed by light-scattering, collected, and stored at −80 °C.

For purification of nuclear capsids, infected HELs were harvested at 48 hpi, cell pellets were washed with 1 × phosphate-buffered saline (PBS) and incubated in NP-40 lysis buffer (0.5% NP-40, 5 M NaCl, 1 M Tris-HCl, pH = 7.0) at 4 °C for 20 min. Nuclei were centrifuged at 2,000 x *g* for 10 min, resuspended in TNE (500 mM NaCl, 10 mM Tris-HCl, 1 mM EDTA, pH = 8.0), sonicated (VCX130, SONICS) at 25% amplitude for 18 s (3s on, 3s off), and centrifuged at 2,000 x *g* for 10 min. This step was repeated once and combined supernatants were centrifuged at 10,000 x *g* for 20 min at 4°C to remove cell debris. Supernatants containing capsids were subjected to iodixanol density gradient centrifugation described above. Capsids were observed as light-scattering bands, collected, and stored at −80 °C.

### Transmission electron microscopy (TEM)

HELFs (Ctl) and the WDR knockdown HELFs (KD) were infected with HCMV at an MOI of 0.5 and harvested at 120 hpi. Cells were fixed with 2.5% (W/V) glutaraldehyde for 1 h at room temperature then treated with 1% osmium tetroxide and dehydrated through a graded series of ethanol concentrations (from 30 to 100%) and embedding with an Embed 812 kit (Electron Microscopy Sciences, Fort Washington, PA). Ultrathin sections (60 to 80 nm) of embedded specimens were prepared and deposited onto formvar-coated copper grids (200-mesh), stained with 2% (W/V) phosphotungstic acid (PTA, pH 6.8), and observed under a Tecnai transmission electron microscope (FEI) operated at 200 kV as described previously (47).

### Immuno-electron microscopy

Purified virions were mock treated or treated with 2% Triton X-100, then adsorbed onto carbon-coated nickel 150 mesh grids for 2 h at room temperature and blocked with 5% goat serum albumin. Grids were incubated with antibodies to WDR5, gB, pp28, or normal mouse IgG overnight at 4 °C in a humidified chamber. Grids were washed three times for 1 min with PBS and then incubated for 30 min at room temperature with 10-nm colloidal gold-affiniPure goat anti-mouse IgG. Grids were washed once with PBS and once with water, then stained with 2% (w/v) phosphotungstic acid for 30 s. Grids were observed using a Tecnai G^2^ 20 TWIN transmission electron microscope (FEI) operated at 200 kV.

### LC-MS/MS analysis

LC-MS/MS analysis was performed on a Q Exactive mass spectrometer (Thermo Scientific) that is coupled to Easy nLC (Proxeon Biosystems, now Thermo Fisher Scientific) for 60 min. The mass spectrometer was operated in positive ion mode. MS data was acquired using a data-dependent top 10 method dynamically choosing the most abundant precursor ions from the survey scan (300– 1800 m/z) for HCD fragmentation. Automatic gain control (AGC) target was set to 3e6, and maximum inject time to 10 ms. Dynamic exclusion duration was 40.0 s. Survey scans were acquired at a resolution of 70,000 at m/z 200 and resolution for HCD spectra was set to 17,500 at m/z 200. Isolation width was 2 m/z. Normalized collision energy was 30 eV and the underfill ratio, which specifies the minimum percentage of the target value likely to be reached at maximum fill time, was defined as 0.1%. The instrument was run with peptide recognition mode enabled.

### Statistical analyses

Data were analyzed by chi-square or one-way ANOVA, as appropriate, using SPSS software (version 18.0; SPSS). Results were shown as means ± one standard deviation from three independent experiments. A value of P < 0.05 was considered significant.

